# Hierarchical design of pseudosymmetric protein nanoparticles

**DOI:** 10.1101/2023.06.16.545393

**Authors:** Quinton M. Dowling, Young-Jun Park, Neil Gerstenmaier, Erin C. Yang, Adam Wargacki, Yang Hsia, Chelsea N. Fries, Rashmi Ravichandran, Carl Walkey, Anika Burrell, David Veesler, David Baker, Neil P. King

## Abstract

Discrete protein assemblies ranging from hundreds of kilodaltons to hundreds of megadaltons in size are a ubiquitous feature of biological systems and perform highly specialized functions ^1–3^. Despite remarkable recent progress in accurately designing new self-assembling proteins, the size and complexity of these assemblies has been limited by a reliance on strict symmetry ^4,5^. Inspired by the pseudosymmetry observed in bacterial microcompartments and viral capsids, we developed a hierarchical computational method for designing large pseudosymmetric self-assembling protein nanomaterials. We computationally designed pseudosymmetric heterooligomeric components and used them to create discrete, cage-like protein assemblies with icosahedral symmetry containing 240, 540, and 960 subunits. At 49, 71, and 96 nm diameter, these nanoparticles are the largest bounded computationally designed protein assemblies generated to date. More broadly, by moving beyond strict symmetry, our work represents an important step towards the accurate design of arbitrary self-assembling nanoscale protein objects.

---

Self-assembling protein complexes are ubiquitous structures foundational to living systems. These structures span length scales from a few nanometers to micron-sized viral capsids and perform a wide variety of structural and biochemical functions ^1–3^. The information driving assembly of these complexes is encoded in their amino acid sequences and functionally takes the form of the structures of individual protein subunits and the interactions between them. The unique properties of self-assembling proteins have been exploited for applications in drug delivery, enzyme encapsulation, and vaccines ^6–9^. However, relying on naturally occurring assemblies constrains the engineer to existing sizes, shapes, and levels of complexity. Methods for generating new self-assembling proteins render additional classes of structures and functions accessible, enabling these properties to be tailored to specific applications ^10^.

Advances in methods for controlling or designing the way protein subunits interact has led to an explosion of new designed assemblies in recent years, particularly those with finite, point-group symmetries (i.e., oligomers, nanoparticles, and capsids) ^11^. Engineered nanoparticles and capsids have been generated by computational protein design ^12–19^, rational design ^20^, genetic fusion and domain swapping ^21–32^, orthogonal chemical interactions such as metal coordination ^33–36^, and laboratory evolution ^37,38^. Each of these methods has a characteristic level of precision and predictive capacity. Computational docking and protein-protein interface design stands out for its ability to consistently create new protein complexes with atomic-level accuracy, although with a relatively modest success rate due to the unique challenge posed by each interface design problem. Nevertheless, computationally designed protein nanoparticles have been engineered to encapsulate small molecules, nucleic acids, and other polymers ^39–41^; evolved for improved cargo packaging and extended *in vivo* half-life ^39^; applied to enhance receptor-mediated signaling and virus neutralization ^15,18,42^; and used as scaffolds for structure determination ^43–45^, multi-enzyme co-localization ^46^, and multivalent antigen presentation^17,47–52^, including in multiple vaccines currently in clinical development ^50,53^ or licensed for use in humans ^54,55^. Further computational methods development will give rise to designed protein nanomaterials of continually increasing sophistication, leading to improved performance in these applications and making additional applications possible.

Design methods reported to date have relied on the use of strict symmetry and pre-existing oligomeric building blocks to reduce the number of new interfaces that must be designed ^4,5^. Although this approach yields access to a handful of finite (i.e., bounded) symmetric architectures that require only a single designed interface ^56^, it nevertheless places a severe constraint on the architectures accessible to design and their size and complexity. The largest and most complex structures designed using this approach comprise 120 subunits and have strict icosahedral symmetry, featuring a single copy of each of two subunits in the icosahedral asymmetric unit ^15–17^. Developing methods for breaking the symmetry of computationally designed protein assemblies is a key next step in developing more sophisticated self-assembling proteins.

Four routes to larger and more sophisticated protein assemblies exist, each of which is observed in naturally occurring self-assembling proteins. First, larger protein subunits could be used as building blocks, with titin providing an extreme example ^57^. However, this approach is untenable as a general solution, as limits in protein translation, folding, stability, and flexibility are quickly encountered ^58^. Second, the number of different kinds of subunits in the assembly (or its asymmetric unit; asu) could be increased by designing new asymmetric interactions between them, as observed in molecular machines such as RNA polymerases ^59^ or NADH-quinone oxidoreductase ^60^. Although ultimately we expect this approach to become possible, it is currently impractical as it would compound the low success rates of existing interface design methods. Third, principles of quasi-equivalence could be used to design large assemblies from protein subunits that adopt subtly different conformations depending on their local symmetry environment, a phenomenon commonly found in icosahedral virus capsids ^37,38,61–63^. However, current computational protein design methods lack the precision required to reliably encode in a single amino acid sequence the multiple subtly different backbone conformations required to implement this approach. Finally, pseudosymmetry could be used to enable asymmetric functionalization of oligomeric building blocks, opening up new routes to the design of larger assemblies. Pseudosymmetry is also frequently observed in icosahedral virus capsids, where genetically distinct subunits or domains adopt roughly symmetric orientations within oligomeric capsomers ^64^. For example, pseudosymmetric trimers in virus capsids may comprise three subunits, each containing two related but slightly distinct domains that result in an (A-B)-(A-B)-(A-B) arrangement with roughly six-fold symmetry at the backbone level ^65–67^. Such trimers can be arranged in hexagonal lattices that form the facets of very large icosahedral assemblies; the A and B domains form the distinct sets of contacts that are necessary to form non-porous lattices. Although designing pseudosymmetric assemblies requires the creation of multiple new protein-protein interfaces, a hierarchical approach in which pseudosymmetric oligomers are designed first and subsequently used as the building blocks of larger pseudosymmetric assemblies would allow the distinct interfaces to be designed and validated individually. This approach avoids compounding the relatively high failure rate of interface design and, as we show, permits the design of novel cage-like protein nanomaterials that far exceed the size and complexity of previously designed assemblies.

## Pseudosymmetric heterotrimer design

We started our pseudosymmetric design with a homotrimeric aldolase from the hyperthermophilic bacterium *Thermotoga maritima* that is remarkably stable and tolerant of modification (PDB ID 1WA3; ^*68*^). This trimer has previously been used to design multiple one- and two-component protein assemblies ^12,16,69^, which as we show below makes possible the re-use of these previously designed interfaces in the creation of large pseudosymmetric assemblies. We set out to identify the minimum set of mutations necessary to drive formation of a pseudosymmetric heterotrimer. We used two methods to identify individual mutations predicted to disrupt—as well as compensatory mutations predicted to restore—homotrimer stability, reasoning that combining sets of such mutations across three variants of the trimer subunit could yield pseudosymmetric heterooligomers (**Fig. 1A**). First, the energetic effects of all possible single and pairwise mutations in 98 contacting residue pairs over 36 positions in the 1WA3 homotrimer interface were evaluated using Rosetta. The Rosetta score and predicted homotrimerization energy (ddG) of each individual and double mutant were normalized to the wild-type (WT) score and ddG by simple subtraction. 96 unique individual mutations had normalized ddG or Rosetta scores >+100, suggesting they may disrupt the wild-type homotrimeric interface (**Fig. S1, A and B**). However, many of these had no compensatory mutation that brought the normalized total score or normalized ddG close to zero. Double mutants that did have favorable normalized score or ddG values were considered further (**red boxes, Fig. 1, B and C**). Second, because 1WA3 is a naturally occurring protein, we also used bioinformatics to guide our mutant and double mutant selection. Using GREMLIN ^70,71^ we inspected the coupling matrices of highly co-evolving residues at the trimer interface to identify low-frequency single mutations (e.g., H91I; PDB ID 1WA3 numbering) with high-frequency compensatory mutations (e.g., V118Y) (**Fig. 1D**). As expected, many of the predicted disrupting mutations identified by computational modeling or bioinformatics were mutations to bulky hydrophobic residues (**Fig. 1E**). Models of those single mutants were visually inspected and then paired with the best-scoring double mutant. In total, mutations from 76 mutant pairs were selected for experimental screening (**Table S1**).

**Fig. 1:**
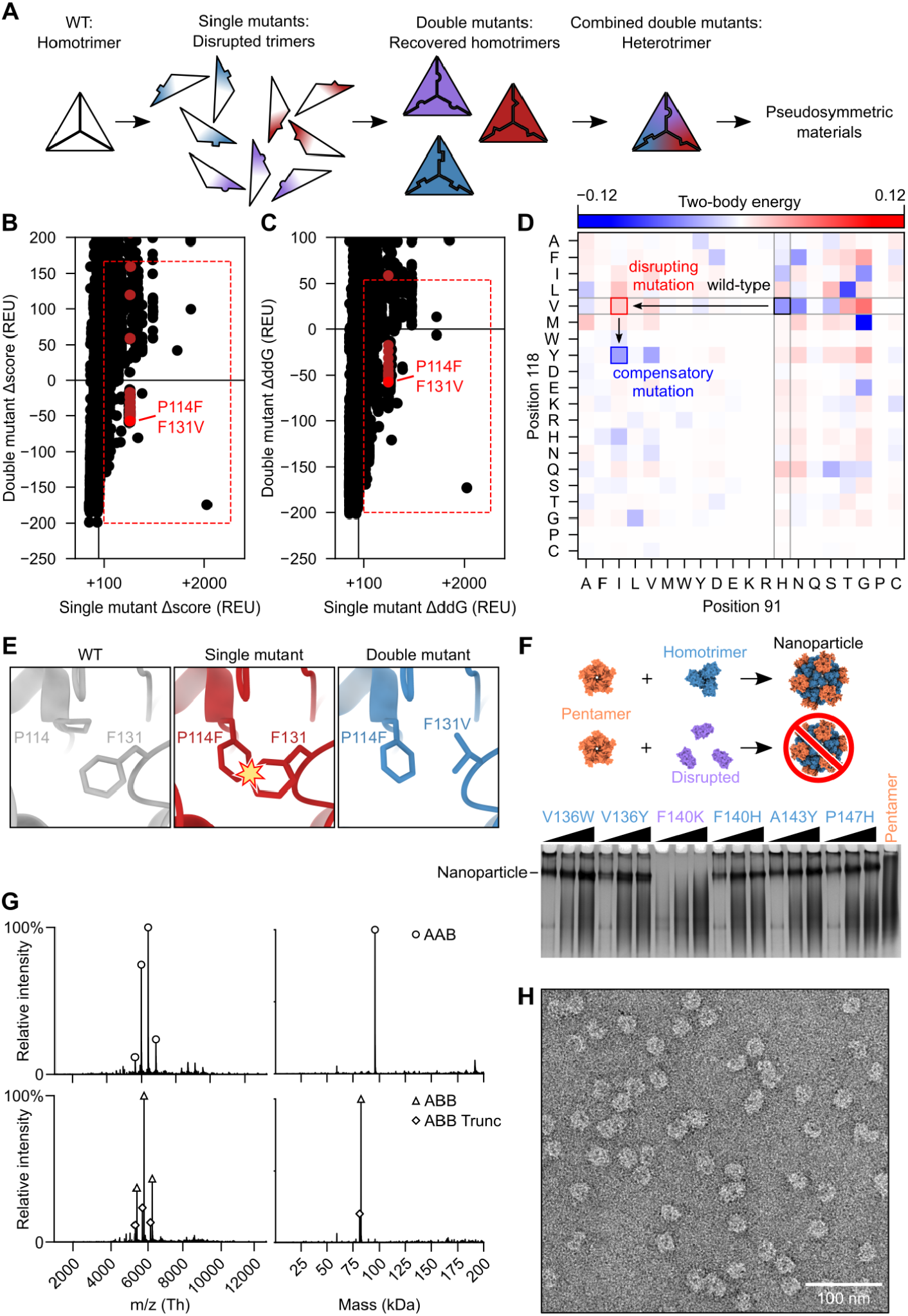
Design and characterization of a pseudosymmetric heterotrimer. (**A**) The design protocol starts with a homotrimer to which single trimer-disrupting mutations are introduced, followed by compensatory mutations that rescue trimer assembly. Sets of orthogonal mutations (depicted as red, purple, and blue) are combined to generate a heterotrimer that can then be used as a component in pseudosymmetric materials. (**B**) Calculated changes in Rosetta score (Δscore) and (**C**) predicted trimerization energy (ΔddG) upon mutation were used to evaluate single mutants (horizontal axis) and double mutants (vertical axis). The red boxes enclose mutants that met selection criteria for further evaluation, and mutant pairs containing P114F, shown in Fig. 1E, are highlighted in red. (**D**) Possible mutations were also evaluated by their co-evolution coupling matrix. Desirable mutations are those where the single mutant/WT pair are observed less often than expected (red; H91I/V118) and the double mutant pair more often than expected (blue; H91I/V118Y). (**E**) An example of a productive mutant pair where the WT residue F131 clashes with the mutant residue P114F and the second mutation F131V resolves the clash. (**F**) Disruption of trimer geometry was assayed by assembling mutant I53-50A trimers in clarified *E. coli* lysates with purified I53-50B pentamer and evaluating the presence or absence of I53-50 nanoparticles by native PAGE. Here, F140K was identified as a disrupting single mutation. Black wedges indicate increasing pentamer concentration in the assembly reaction. (**G**) Native mass spectrometry of (*top*) AAB-enriched and (*bottom*) ABB-enriched heterotrimer fractions purified by IMAC and SEC. ABB Trunc refers to a truncation product of the A chain in which the N-terminal 10 residues of the protein were missing. (**H**) Assembly of I53-50-like nanoparticles using an AAB/AAB mixture of I53-50A heterotrimer was verified by negative-stain EM.

First, the ability of each single mutation to disrupt trimer formation was screened in a lysate-based assay by evaluating its effect on the assembly of I53-50, a previously reported two-component nanoparticle comprising a trimeric component (I53-50A) derived from the 1WA3 trimer ^16^. Clarified lysates from *E. coli* expressing the I53-50A mutants were mixed with purified I53-50B pentamer at three different pentamer concentrations and analyzed by native (i.e., non-denaturing) PAGE (**Fig. 1F**). Four of the 82 single mutants tested failed to yield a band corresponding to the assembled I53-50 nanoparticle, indicating that these either prevented soluble expression of the trimer variant or altered its geometry so that it was no longer assembly-competent. Mutations that prevented nanoparticle formation were then combined with their compensatory mutation to determine if the combination restored the ability to form I53-50 nanoparticles (**Fig. S2**). Through these analyses, three pairs of disrupting and compensatory mutations were identified: H91I/V118Y, P90F/P147A, and P114F/F131V.

We initially set out to generate an “ABC” heterotrimer, in which each subunit has a different amino acid sequence, by combining the three mutant pairs in a tricistronic expression construct (all novel amino acid sequences provided in **Table S2**). The A chain contained the V118Y mutation, the B chain contained F131V and P90F, and the C chain contained H91I, P114F and P147A. However, when we purified the expressed proteins by immobilized metal affinity chromatography (IMAC) and StrepTrap chromatography, analysis by SDS-PAGE suggested the presence of trimers comprising predominantly a mixture of the A and B subunits, with little C subunit (**Fig. S3, A and B**). We therefore expressed a bicistronic “AB” gene, which resulted in a mixture of two distinct heterooligomers we separated by IMAC and size exclusion chromatography (SEC) and identified by native mass spectrometry: a trimer comprising one copy of the A chain and two copies of the B chain (“ABB”), as well as a trimer comprising two copies of the A chain and one copy of the B chain (“AAB”) (**Fig. 1G** and **Fig. S3C**). As we describe below, such heterooligomers provide a simple route to designing large pseudosymmetric materials. To confirm that symmetry was preserved at the backbone level—a prerequisite for our hierarchical design approach—we determined if the heterotrimer mixture was assembly-competent by purifying and incubating it in a 1:1 molar ratio with I53-50B pentamer. Assemblies were purified by SEC and the presence of nanoparticles with the known I53-50 morphology ^16^ were observed by negative stain electron microscopy (nsEM;**Fig. 1H**).

### Design and characterization of a 240-subunit, T=4 nanoparticle

We then used the pseudosymmetric heterotrimers to design large, pseudosymmetric assemblies with icosahedral symmetry. We had previously used the 1WA3 homotrimer to generate a single-component nanoparticle with icosahedral symmetry, I3-01, by designing a novel protein-protein interface with two-fold symmetry between the subunits of adjacent trimers ^12^. The existence of this interface allowed us to generate a fifteen-subunit “pentasymmetron” ^72^ comprising five trimers by simply including the I3-01 mutations on the A chains of the AAB heterotrimer (**Fig. 2A**). Docking this pentasymmetron against C3-symmetric homotrimers (“CCC”) and designing novel sequences that create favorable interfaces between the B and C chains yielded models of 240-subunit nanoparticles with icosahedral symmetry (**Fig. 2, B-D**). The Caspar-Klug triangulation number notation ^61^ is useful for describing these pseudosymmetric nanoparticles, although the assignment of subunits to geometric elements is different than in traditional use of the T number in structural virology. In our pseudosymmetric nanoparticles, trimeric building blocks form wireframe-like structures surrounding roughly pentagonal and hexagonal pores, each subunit interacting with exactly one other subunit from a different trimer. The original I3-01 nanoparticle can be thought of as T=1, with one (A) subunit in the asu, while the pentasymmetron-containing 240-subunit nanoparticles are T=4, with four (2×A, 1×B, 1×C) subunits in the asu. In these assemblies *k*=0, so the T number is equal to *h*^*2*^, where *h* is a positive integer representing the number of steps required to traverse from one pentasymmetron to another, each step moving to the next pentagonal or hexagonal pore. Because this is one of the set of equations used to define class I Goldberg polyhedra ^73^, we refer to these nanoparticles using the naming convention GI_T_-X, where G stands for Goldberg, I for icosahedral symmetry, T is used to denote the triangulation number of a particular architecture, and X is a unique identifier for each design. We expressed three initial designs in *E. coli* as tricistronic genes with a 6×His tag on only the C chain, and found that Ni^2+^ beads co-precipitated all three subunits of two of the designs, suggesting assembly (**Fig. S4**). We moved forward with the better expressing and more soluble of the two, GI_4_-F7. To scale up expression of the AAB heterotrimer so that we could explore assembly of GI_4_-F7 *in vitro* from purified components ^74^, we re-cloned it as a bicistronic AB construct with a 6×His tag on the A chain. Upon gradient elution during IMAC, we observed three peaks corresponding to an ABB-rich fraction, an AAB-rich fraction, and off-target AAA homotrimers that assembled to an I3-01-like nanoparticle (**Fig. S5A**). We polished the AAB peak by size exclusion chromatography (SEC), discarding the I3-01-like nanoparticle fraction (**Fig. S5B-D**). In parallel, we purified 6×His-tagged CCC homotrimer—which was also derived from the 1WA3 trimer—by IMAC and SEC.

**Fig. 2.**
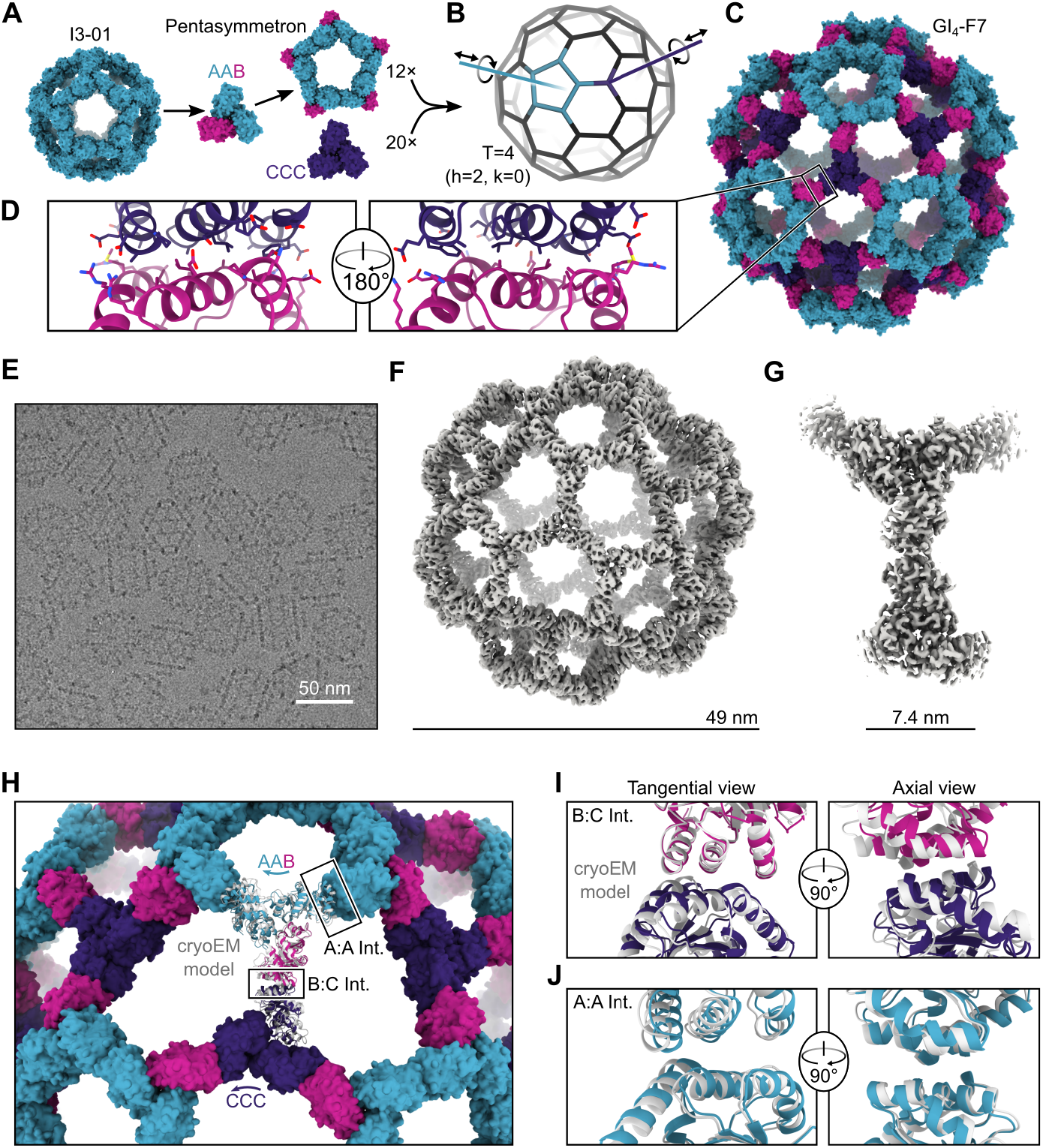
Design and characterization of the 240-subunit GI_4_-F7 nanoparticle. (**A**) Schematic of pentasymmetron generation from I3-01 and the AAB heterotrimer. The A (cyan) subunits in the pentasymmetron retain the two-fold symmetric I3-01 nanoparticle interface, while the B (magenta) subunits are available for docking. (**B**) Docking the pentasymmetron as a rigid body against CCC homotrimers (purple) yields 240-subunit, T=4 assemblies. Translational and rotational DOFs for the pentasymmetron and homotrimer components are indicated. (**C**) A design model of GI_4_-F7. (**D**) Detail of the computationally designed interface between the B and C subunits of GI_4_-F7 design model. (**E**) CryoEM micrograph of assembled GI_4_-F7 nanoparticles embedded in vitreous ice. (**F**) 4.4 Å resolution density map of the entire GI_4_-F7 nanoparticle. (**G**) 3.1 Å resolution density map from an asymmetric unit obtained via symmetry-expansion and local refinement. (**H**) Comparison of the cryoEM structure derived from local refinement (gray ribbon) with the computational design model (colored ribbons), aligned using a single copy of the asu. Arrows indicate rigid body deviations of the pentasymmetron (cyan) and CCC homotrimer (purple). (**I**) Detail of the rigid body deviations from the design model at the B-C interface and (**J**) the A-A (I3-01) interface. In panel J, two neighboring copies of the AAB heterotrimer from the full nanoparticle reconstruction and the design model were aligned.

We mixed the AAB heterotrimer with an excess of the CCC homotrimer in the presence of detergent and initiated assembly by dialyzing overnight into Tris-buffered saline (TBS). The major assembly product was purified by SEC (**Fig. S6**), and images obtained by cryo-electron microscopy (cryoEM) of vitrified specimens revealed wireframe structures with large hexagonal pores that strikingly resembled the design model (**Fig. 2E**). We determined a single-particle reconstruction of GI_4_-F7 at 4.4 Å resolution applying icosahedral symmetry and a 3.1 Å resolution structure of the four chains of the asu (cryoEM processing details in**Fig. S7**and**Table S3**). The cryoEM structure agrees well with the design model, with a Cαroot mean square deviation (RMSD) of 9.3Å across all 240 subunits and 3.0 Å within the asu (**Fig. 2F and G**, **Fig. S8**). The differences between the cryoEM structure and design model are mostly accounted for by slight rigid body deviations allowed by the limited degrees of freedom (DoFs) of the oligomeric building blocks in this symmetric architecture (**Fig. S8A**). The main rigid body deviation is a 5.9° clockwise rotation of the pentasymmetron, accompanied by a 5.8 Å translation away from the origin (**Fig. 2H**). The CCC homotrimer compensates by rotating 12.4° and translating 4.0 Å, resulting in only slight local shifts relative to the design model (2.1 Åacross the B:C subunits;**Fig. 2I**). Within the pentasymmetron, the DoFs of the AAB heterotrimer are no longer restricted by the strict icosahedral symmetry of I3-01, resulting in a slight deviation from perfect two-fold symmetry between neighboring A chains (1.4 ÅCαRMSD; **Fig. 2J** and **Table S4**). The effects of these deviations on the atomic contacts at the A:A, A:B, and B:C interfaces are highlighted in **Fig. S8**. There were minor structural deviations within each subunit, primarily in the B:C interface (**Fig. S9**). Overall, the diameter of GI_4_-F7 observed by cryoEM is within 2% of the design model, establishing that our method is capable of accurately designing pseudosymmetric protein nanomaterials comprising hundreds of subunits.

### Observation and structure of a 540-subunit, T=9 nanoparticle

Unexpectedly, in a number of the GI_4_-F7 micrographs we also observed a 71 nm nanoparticle with a similar wireframe morphology and hexagonal pores (**Fig. 3A**). By counting the hexagonal pores we found that *h*=3; thus the nanoparticle is T=9 and we refer to it as GI_9_-F7. GI_9_-F7 can be explained by the presence of small amounts of ABB heterotrimer in AAB heterotrimer preparations. Analogous to the AAB heterotrimer, which forms a pentasymmetron through five roughly two-fold-symmetric A:A interfaces inherited from I3-01, the ABB heterotrimer forms a two-trimer “disymmetron” structure held together by the same A:A interaction (**Fig. 3B**). In GI_9_-F7 this disymmetron occupies the icosahedral two-fold symmetry axes, providing the edges that connect three-fold-symmetric facets containing three ABB heterotrimers and three CCC homotrimers. As a result, GI_9_-F7 is quasisymmetric in addition to being pseudosymmetric: the A, B, and C subunits each occupy multiple, distinct environments in the assembly. We expanded GI_4_-F7 to generate a design model for GI_9_-F7 containing 12 pentasymmetrons constructed from AAB heterotrimers, 30 disymmetrons comprising ABB heterotrimers, and 60 CCC homotrimers (**Fig. 3B**). The asu of GI_9_-F7 therefore comprises one AAB trimer, one ABB trimer, and one CCC trimer. To generate more homogenous preparations of GI_9_-F7, we separately polished the AAB and ABB heterotrimer fractions from IMAC (see **Methods**) and assembled them with CCC homotrimer at a 1:1:1 ratio. Micrographs of SEC-purified GI_9_-F7 assemblies revealed enrichment of the target assembly (**Fig. S10**), and we determined a cryo-EM structure of the nanoparticle to 6.7 Å resolution applying icosahedral symmetry, as well as a 4.0 Å resolution structure of the asu (**Fig. 3C, Fig. S7**, and **Table S3**). Consistent with the accuracy with which we designed GI_4_-F7, the GI_9_-F7 cryoEM structure deviates from the design model by only 11.5 Å Cα RMSD across all 540 subunits, or 1.6% of the nanoparticle diameter, and superimposition of the designed asu with the structure yields a Cα RMSD of 4.6 Å across all nine chains.

**Fig. 3.**
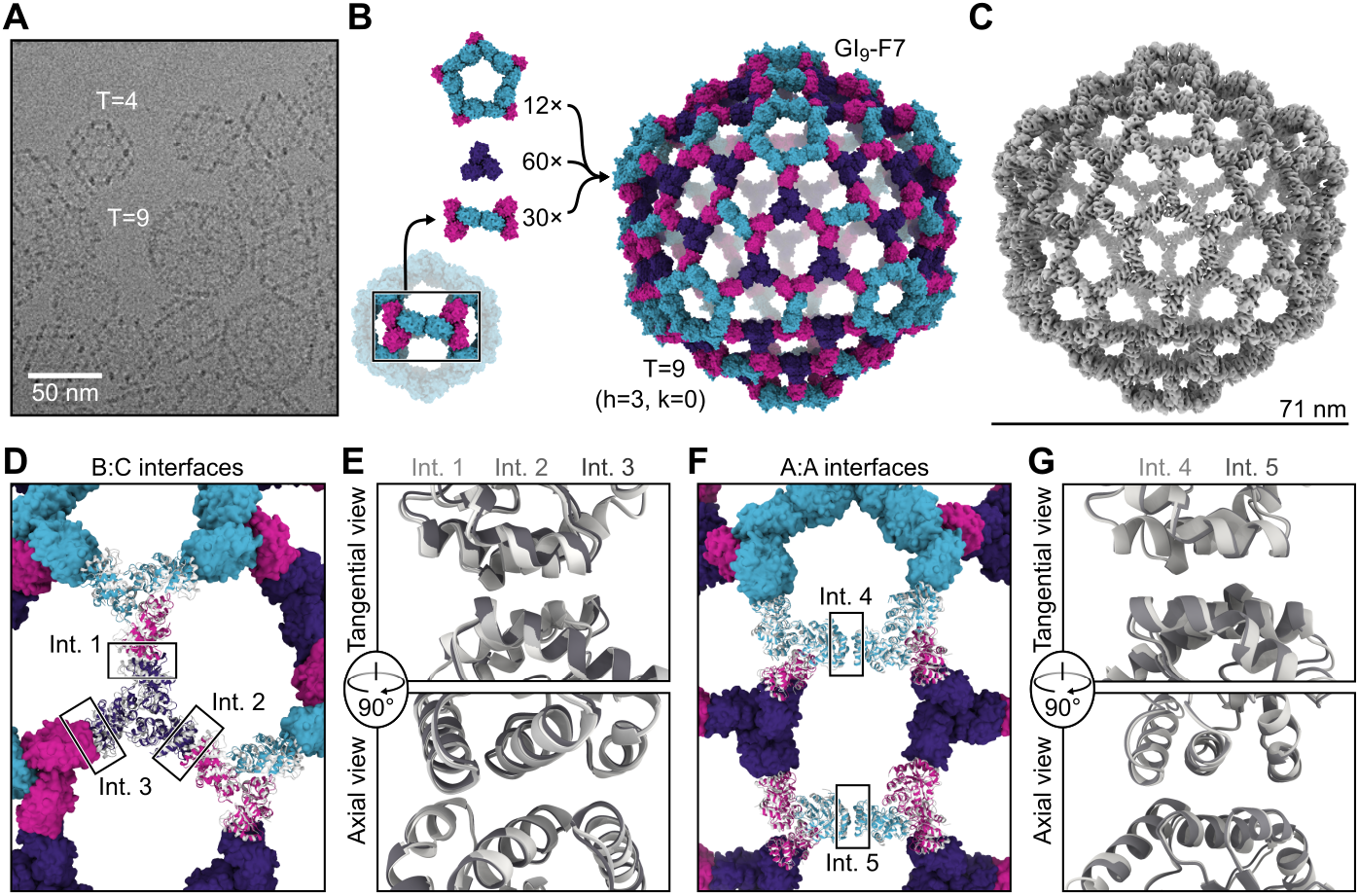
Discovery and characterization of the 540-subunit GI_9_-F7 nanoparticle. (**A**) A cryo-electron micrograph showing GI_4_-F7 and GI_9_-F7 nanoparticles in the same preparation. (**B**) GI_9_-F7 comprises, from top to bottom, 12 pentasymmetrons consisting of five copies of the AAB heterotrimer, 60 copies of the CCC homotrimer, and 30 copies of the disymmetron consisting of two copies of the ABB heterotrimer. A subunits are cyan, B subunits are magenta, and C subunits are purple. (**C**) CryoEM map of GI_9_-F7 at 6.7 Å resolution. (**D**) Comparison of a model derived from the cryoEM map (gray) with the computational design model (colors), aligned using a single asu (shown in cartoon). The three independent copies of the B:C interface in the asu are indicated. (**E**) Alignment of Int. 1. (light gray), Int. 2 (medium gray), and Int. 3 (dark gray) from the cryoEM model. (**F**) Alignment of two neighboring copies of AAB heterotrimers from the cryoEM model to the design model. The two independent copies of the A:A (I3-01) interface, located in the pentasymmetron (Int. 4) and the disymmetron (Int. 5), are indicated. (**G**) Alignment of the two A:A (I3-01) interfaces from the cryoEM model, Int. 4 (light gray) and Int. 5 (medium gray).

Although the pentasymmetron, disymmetron, and three homotrimers in GI_9_-F7 are respectively constrained by the five-fold, two-fold, and three-fold icosahedral symmetry axes, no single trimer occupies a position constrained by icosahedral symmetry—the icosahedral three-fold instead passes through a large pore. Each trimer can therefore deviate from the design model along all six rigid body DoFs. As a result, the two designed nanoparticle interfaces (B:C and A:A) occupy five quasi-equivalent positions in GI_9_-F7. Two of the B:C interfaces are located within the icosahedral asu, between the CCC homotrimer and the B chain of a neighboring pentasymmetron (interface 1) or disymmetron (interface 2) (**Fig. 3D**). The third B:C interface is between the CCC homotrimer and the B chain of a disymmetron in a neighboring asu (interface 3). Despite being unconstrained by symmetry, interfaces 1-3 fit well to the density, with a very small deviation from the design model comprising only a small rotation with very little radial translation (**Fig. 3E, Fig. S11A-D**, and **Table S5**). Interfaces 4 and 5 are the A:A (i.e., I3-01) interfaces in the pentasymmetron and disymmetron, respectively (**Fig. 3F**). Unlike interfaces 1-3, interfaces 4 and 5 appear to differ, with a Cα RMSD of 1.3 Å to each other and Cα RMSDs of 2.0 and 2.4 Å to the GI_9_-F7 design model, respectively (**Fig. 3G** and **Fig. S11E-G**). This difference arises because the pentasymmetron interface (interface 4) is not symmetrically constrained, while the disymmetron interface (interface 5) is constrained by the icosahedral two-fold symmetry axis. We propose that the lowest-energy state of the A:A interface is not perfectly symmetric, but that the symmetry requirements for nanoparticle assembly force it to adopt a higher-energy, two-fold-symmetric configuration where appropriate.

### Generation of pseudosymmetric nanoparticles with extensible hexagonal lattice facets

The geometries of I3-01, GI_4_-F7, and GI_9_-F7 are analogous to the first three instances in the infinite series of class I Goldberg polyhedra ^61,73,75^. The larger instances in this series are effectively constructed by folding 20 roughly triangular two-dimensional (2D) hexagonal lattices into icosahedron-like shapes through the introduction of curvature at their edges and vertices. Theoretically, the next nanoparticle in the series would be GI_16_-F7. As in GI_4_-F7 and GI_9_-F7, curvature in this structure would be provided by disymmetrons and pentasymmetrons. However, GI_16_-F7 would have a C3-symmetric component centered on the icosahedral three-fold symmetry axis, as opposed to the pore-centered three-fold of GI_9_-F7. Extrapolating from GI_4_-F7 and GI_9_-F7, this component must be a homotrimer of the B chain (“BBB”), and it must be coplanar with the six surrounding CCC homotrimers (i.e., their three-fold axes must be parallel; **Fig. 4A**). Nanoparticles beyond GI_16_-F7 simply add more copies of the BBB and CCC homotrimers (and ABB disymmetrons) to form larger two-dimensional hexagonal arrays (**Fig. 4B**). Thus, obtaining GI_16_-F7 and the larger nanoparticles in the series does not require new interface design, only production of BBB homotrimer. Indeed, analyzing an equimolar mixture of purified BBB and CCC homotrimers by nsEM yielded a 2D array with a characteristic hexagonal lattice diffraction pattern (**Fig. S12A-C**). The dimensions of the array agree well with a design model derived from the GI_9_-F7 nanoparticle (**Fig. S12, D and E**).

**Fig. 4.**
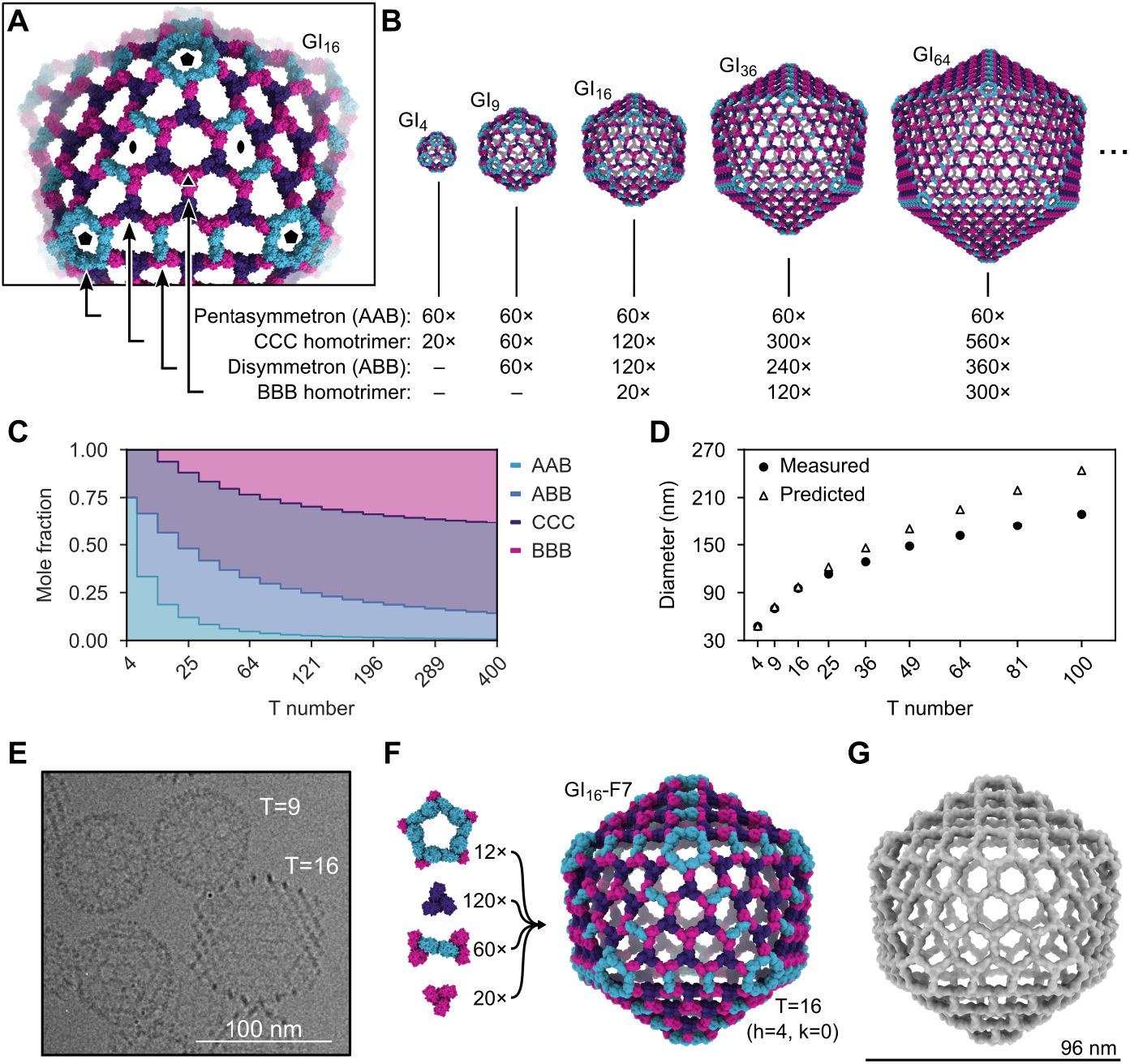
Generation of pseudosymmetric nanoparticles with extensible hexagonal lattice facets. (**A**) The four types of trimers required to generate pseudosymmetric nanoparticles with T numbers ≥16, viewed in the context of the GI_16_-F7 design model. Icosahedral five-, two-, and three-fold symmetry axes are indicated. (**B**) Design models and corresponding building block stoichiometries of GI_4_-F7, GI_9_-F7, GI_16_-F7, GI_36_-F7, and GI_64_-F7 nanoparticles. The stoichiometries listed indicate the number of each kind of trimeric component. (**C**) Graphical depiction of trimer stoichiometry as a function of T number. (**D**) Theoretical nanoparticle diameters and Z-average hydrodynamic diameters measured by DLS as a function of trimer stoichiometry used during *in vitro* assembly (indicated by T number). (**E**) Cryo-electron micrograph of a sample containing GI_9_-F7 and GI_16_-F7 nanoparticles. (**F**) Composition and design model of the GI_16_-F7 nanoparticle. (**G**) 14.9 Å resolution cryoEM map of GI_16_-F7.

In the absence of other control mechanisms, the inclusion of BBB homotrimer in assembly reactions should yield distributions of large T-number assemblies rather than monodisperse preparations of a single species. However, the relative stoichiometries of these components in each assembly vary as a function of T number (**Fig. 4, B and C**), providing a potential mechanism for controlling assembly size. We prepared assembly reactions containing the four components at the stoichiometries corresponding to T=4, 9, 16, 25, 36, 49, 64, 81, and 100 nanoparticles. Consistent with our predictions, the Z-average hydrodynamic diameter measured by DLS increased with increasing target T number (from 47.5 ± 0.4 nm to 188 ± 1.1 nm), though the observed hydrodynamic diameter deviated from the predicted diameter at higher T numbers (**Fig. 4D**,**Table S6**). GI _16_-F7 was readily observed by cryoEM in assembly reactions prepared at the T=16 stoichiometry (**Fig. 4E**). GI_16_-F7 is predicted to have a diameter of 96 nm and contains 12 pentasymmetrons, 120 CCC homotrimers, 60 disymmetrons, and 20 BBB homotrimers for a total of 960 subunits (**Fig. 4F**). We determined a 14.9 Å resolution cryoEM map of GI_16_-F7 and found that it closely matches the expected geometry of the design (**Fig. 4G**). This assembly has an internal volume that is roughly 90-fold larger than our previously designed nanoparticles with strict icosahedral symmetry ^12,16^ and adeno-associated viruses, commonly used vectors for gene therapy ^76^.

## Conclusions

Here we show that designing pseudosymmetric protein building blocks, in which symmetry is broken at the sequence level while backbone symmetry is maintained, enables the construction of very large pseudosymmetric protein nanoparticles. By breaking the reliance on strict symmetry, this work represents an important leap forward relative to established methods for accurately designing novel self-assembling proteins ^13,14,16,17,24,31,32,77^. Although those methods are general with respect to the choice of building block and can therefore give rise to a rich variety of potential assemblies, the space of strictly symmetric structures is nevertheless infinitesimally small compared to the space of asymmetric architectures. Continued development of computational methods for designing new proteins and protein assemblies, docking existing protein building blocks, and designing protein-protein interfaces will enable the generation of increasingly asymmetric protein nanomaterials. The technological impact of such computationally designed protein nanomaterials is highlighted by the recent regulatory approval and commercialization of a designed protein nanoparticle vaccine for COVID-19 ^54^.

Although some small viruses make purely pseudosymmetric capsids, many larger capsids are constructed by combining pseudosymmetry with quasisymmetry. Analogously, while GI_4_-F7 and the assemblies reported in the accompanying manuscript (Lee & Kibler, et al.) are pseudosymmetric, with each distinct subunit in a single chemical environment, GI_9_-F7 and larger structures are also quasisymmetric, with genetically identical subunits in more than one chemical environment. The A subunit occupies an asymmetric position in the pentasymmetron and either an asymmetric position in the disymmetron (for even T numbers) or both asymmetric and two-fold symmetric positions in disymmetrons (for odd T numbers >9). Likewise, the B and C subunits occupy different chemical environments depending on their locations in the assembly. Quasisymmetry is enabled by the use of two-component heterotrimers (ABB and AAB), which provides for economy in coding for larger assemblies. The T=4 structure requires only three distinct chains, compared to four chains for the more conceptually straightforward approach of a strictly pseudosymmetric ABC heterotrimer and DDD homotrimer. For larger particles the economy is even greater: for example, only three unique chains are required to make T=9 nanoparticles, but seven would be needed for the strictly pseudosymmetric approach. The tradeoff to this economy is a reduction in precision compared to the strictly pseudosymmetric approach we describe in the accompanying manuscript (Lee & Kibler, et al.), although as we have shown this can be partially overcome by controlling the stoichiometry of the assembly reaction.

We used a hierarchical design strategy to fulfill the requirement for multiple designed interfaces in our pseudosymmetric nanomaterials. By constructing pseudosymmetric heterotrimers first and combining these with an existing designed interface to generate pentasymmetrons, we produced 240-subunit and larger pseudosymmetric assemblies in a single dock-and-design step. Similar hierarchical and modular design strategies are widespread in reticular chemistry ^78,79^ and DNA nanotechnology ^80–83^, and should become increasingly powerful in protein nanomaterials design as the number and kinds of modular protein building blocks continue to increase ^84–89^.

## Acknowledgements

The authors thank Samuel Wrenn and the IPD Nanoparticle Core for assisting with protein production and purification, Florian Praetorius and the King laboratory for comments on the manuscript, and Ratika Krishnamurty for program management. Native mass spectrometry measurements were provided by the Wysocki lab at the Ohio State University. pCDB179 was a gift from Christopher Bahl (Addgene plasmid #91960).

## Data and materials availability

Further information and requests for resources and reagents should be directed to and will be fulfilled by the corresponding author. All data are available in the manuscript or the supplementary materials. EM maps and models of structures will be deposited to the EMDB and PDB prior to publication.

## Author Contributions

Q.M.D., Y.H., D.B., N.P.K., conceived the study; Q.M.D. designed the nanomaterials; Q.M.D. performed bioinformatics analyses; Q.M.D., N.G., and A.B. produced and experimentally characterized pseudosymmetric mutants; Q.M.D, N.G,. and R.R., developed purification methods used in this study; C.W. developed the 12 negative mutants used to facilitate production of the pentasymmetron; Q.M.D., N.G., and R.R., produced and characterized the nanoparticles. Q.M.D., N.G., E.Y., A.W., Y.H., and N.P.K., determined the geometric principles of assembly; Q.M.D. and C.F. produced, characterized, and collected nsEM data of the 2D-arrays; Y.J.P collected the CryoEM data. Y.J.P. and D.V.. analyzed and processed the CryoEM data; all authors analyzed data; Q.M.D., and N.P.K. wrote and revised the manuscript.

## Methods

### Pseudosymmetric trimer design

To identify mutations for altering trimer assembly specificity, we first identified all pairs of interacting residues in the trimer interface. Contacts were defined as any residue with a heavy (i.e., non-hydrogen) atom within 4 Åof a heavy atom in a residue across the interface. We then used Rosetta to calculate the total score of poses containing all possible pairs of mutations, as well as the difference in score between the trimeric and monomeric states using the ddG filter. Individual mutations were evaluated by comparing their ddG and total scores to those of the WT interface according to Equation 1. The total scores and ddG values of the paired mutations were similarly normalized according to Equation 2.

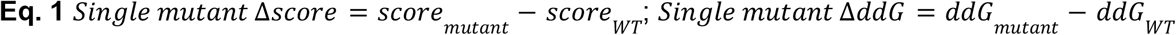

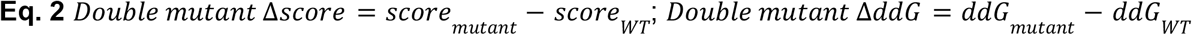

Ideal mutant pairs were those where one or both single mutations increased the energy of the trimer relative to the WT (i.e., normalized scores > 0) while the double mutation had no effect or stabilized the trimer (i.e., normalized scores ≤0). We also identified likely positions for design using coevolutionary analysis ^70,71^. Strongly co-evolving residues at the protein-protein interface were identified using GREMLIN. We then identified mutations that were negatively correlated with the WT pair for testing experimentally.

### Mutant protein expression

Mutant I53-50A trimers were expressed at three scales. Small-scale expressions were performed at 1 mL culture volume in 96-well plates with 2 mL well volume. Medium-scale expressions were performed at 50 mL culture volume in 250 mL baffled shake flasks. Large-scale expressions were performed at 500 mL culture volumes in 2 L baffled shake flasks. All proteins were expressed in T7 competent *E. coli* in TB media, with IPTG induction for 3 hours at 37 °C. Cells were pelleted and frozen at -20 °C until lysis. Prior to lysis cells were defrosted on ice in lysis buffer (50 mM Tris pH 8.0, 250 mM NaCl, 20 mM imidazole, 1 mM phenylmethylsulfonyl fluoride (PMSF), 1 mM dithiothreitol (DTT), 0.1 mg/mL DNase, and 0.1μM RNase, unless otherwise noted). Small-scale expressions were lysed with a plate sonicator (QSonica), medium-scale expressions were lysed with a probe sonicator, and large-scale expressions by microfluidization (18,000 psi, one pass). Lysates from small-scale expressions were clarified by centrifugation in a swinging bucket rotor at 4,000 rcf. Lysates from medium- and large-scale expression lysates were clarified by centrifugation at 12,000 rcf in a fixed-angle rotor.

### I53-50B expression and purification

Pentameric I53-50B was produced recombinantly in *E. coli*. A pET29b expression plasmid encoding I53-50B.4PT1 ^16^ was synthesized by GenScript using the NdeI and XhoI restriction sites with a double stop codon just before the C-terminal polyhistidine tag. Tagless protein was expressed in Lemo21(DE3) cells (NEB) in LB (10 g Tryptone, 5 g Yeast Extract, 10 g NaCl) grown in a 10 L BioFlo 320 Fermenter (Eppendorf). At inoculation, impeller speed was set to 225 rpm, SPLM set to 5 with O_2_ supplementation as part of the dissolved-oxygen aeration cascade, and the temperature set to 37 °C. At the onset of a DO spike (OD ∼12), the culture was fed with a bolus addition of 100 mL of 100% glycerol and induced with 1 mM IPTG. During this time, the culture temperature was reduced to 18 °C and O_2_ supplementation was ceased, with expression continuing until OD reached ∼20. The culture was harvested by centrifugation and the protein was purified from inclusion bodies. First, pellets were resuspended in PBS, homogenized, and then lysed by microfluidization using a Microfluidics M110P at 18,000 psi. Following sample clarification by centrifugation (24,000 g for 30 min), the supernatant was discarded and protein was extracted from the pellet using a series of three washes. The first wash consisted of PBS, 0.1% Triton X-100, pH 8.0. The second wash consisted of PBS, 1 M NaCl, pH 8.0, and the final wash (extraction) consisted of PBS, 2 M urea, 0.75% CHAPS (3-[(3-Cholamidopropyl)dimethylammonio]-1-propanesulfonate), pH 8.0. Following extraction, the sample was applied to a DEAE Sepharose FF column (Cytiva) on an AKTA Avant150 FPLC system (Cytiva). After sample binding, the column was washed with 5 column volumes of PBS at pH 8.0 with 0.1% Triton X-100, followed by a wash with 5 column volumes of PBS at pH 8.0 with 0.75% CHAPS. The protein was eluted with 3 column volumes of PBS at pH 8.0 with 500 mM NaCl. After purification, fractions were pooled and concentrated in 10K MWCO centrifugal filters (Millipore), sterile filtered (0.22μm), aliquoted and flash frozen in liquid nitrogen, and stored at -80 °C until use.

### Assembly competency analysis

Single mutations were introduced into the I53-50A trimer ^16^ by QuikChange™ site-directed mutagenesis. Sequence-verified mutants were expressed at small scale (see above). Clarified lysates were separated from pellets and a 5 μL aliquot set aside for characterization by SDS-PAGE. The pellet was resuspended in lysis buffer and a 5 μL aliquot set aside for characterization by SDS-PAGE. Clarified lysate was immediately mixed with purified I53-50B.4PT1 pentamer. Because trimer expression levels varied from mutant to mutant, pentamer was added at three different concentrations. To 10 μL lysate, 7.5, 2.5, or 0 μL lysis buffer was added, followed by 2.5, 7.5, or 10 μL I53-50B.4PT1 pentamer at 1.8 mg/mL. The assembly reaction was allowed to proceed for 30 minutes at room temperature. A control assembly of previously purified I53-50A trimer and I53-50B.4PT1 pentamer at a 1:1 molar ratio was performed in parallel and included on all native PAGE gels. A 10 μL aliquot of each assembly reaction was mixed 1:1 with Native Sample Buffer (Bio-Rad Laboratories), loaded into precast 4-15% polyacrylamide gels (Bio-Rad Laboratories), and run with 1× Tris-Glycine Native PAGE buffer for 3 hours at 200 V. The gel was stained with GelCode Blue™ (Thermo Fisher Scientific) and destained in water. The lack of an I53-50 nanoparticle band on the native gel indicated single mutations that disrupted either trimer formation or trimer geometry such that the mutant trimer was no longer assembly competent.

### Screening of mutant combinations

Single mutants that disrupted I53-50A trimer—and therefore I53-50 nanoparticle—formation were then combined with “rescue” mutations intended to generate pseudosymmetric I53-50A trimers. Synthetic DNA encoding potential combinations were ordered as heterotrimeric operons cloned into pCDB179 from IDT. To facilitate detection of the distinct components of the heterotrimer, a 6×His-SUMO domain was added to one subunit and sfGFP and an avi-tag added to a second subunit via genetic fusion. The third subunit bore a Strep-tag via genetic fusion. Variants were tested for I53-50 nanoparticle formation using trimer-containing *E. coli* lysates and purified I53-50B pentamer as described above. Combinations that formed I53-50 nanoparticles were expressed at large scale (see above) and purified by Ni^2+^ affinity chromatography on a HisTrap FF column (Cytiva). Briefly, clarified lysate was passed through a pre-equilibrated 5 mL HisTrap FF column, washed with 3-5 column volumes of wash buffer (50 mM Tris pH 8.0, 250 mM NaCl, 20 mM imidazole, 1 mM DTT), and heterotrimer was eluted with either a step elution or a gradient over 40 minutes at 3 mL/min flow rate into 100% elution buffer (50 mM Tris pH 8.0, 250 mM NaCl, 500 mM imidazole, 1 mM DTT). Major fractions corresponding to the two observed peaks in the elution profile were pooled seperately, concentrated in a 30 kDa cutoff Amicon concentrator (Millipore), and injected onto a pre-equilibrated Superdex 200 Increase 10/300 column (Cytiva). The SEC buffer was 25 mM Tris pH 8.0, 150 mM NaCl, 1 mM DTT. Fractions corresponding to the trimer peak from each chromatogram were collected for analysis by native mass spectrometry (MS). Alternatively the IMAC eluate was pooled and loaded onto a StrepTrap HP column (Cytiva) pre-equilibrated in binding buffer (100 mM Tris pH 8.0, 150 mM NaCl, 1 mM EDTA, and 1 mM DTT). The column was then washed with 10 column volumes of binding buffer, or until the A280 absorbance leveled off at baseline and eluted with a step elution in binding buffer plus 2.5 mM desthiobiotin. Major fractions were analyzed by reducing SDS-PAGE. Westerns were performed on select fractions with either an anti-Avi or anti-Strep primary antibody.

### Native mass spectrometry

Trimer purity, identity, and oligomeric state were analyzed by on-line buffer exchange MS ^90^ in 200 mM ammonium acetate using a Vanish ultra-performance liquid chromatography coupled to a Q Exactive ultra-high mass range Orbitrap mass spectrometer (Thermo Fisher Scientific). The recorded mass spectra were deconvolved with UniDec version 4.2+ ^91^.

### Assembly of I53-50 nanoparticles using pseudosymmetric I53-50A heterotrimer

The native MS-verified pseudosymmetric I53-50A heterotrimer was expressed and purified at medium scale as described above and mixed at a 1:1 molar ratio with purified I53-50B.4PT1 pentamer and allowed to assemble at room temperature for 30 minutes. Assembled nanoparticles were characterized by DLS and negative-stain EM as described below.

### Computational design of T=4 nanoparticles

We created a model of the pentasymmetron by extracting five trimers surrounding the icosahedral five-fold from I3-01 ^12^. We reverted the interface residues on the unpaired subunit back to the original 1WA3 sequence, mutated 12 residues to negatively charged amino acids to enhance expression and facilitate purification, then combined each trimer into a single chain so that the pentasymmetron could be treated computationally as a simple homopentamer. We used previously described protocols ^13,16^ to dock and design T=4 nanoparticles. Docked configurations were manually screened to ensure interfaces were between the unpaired pentasymmetron subunit and the homotrimer. Designs were visually inspected and any overly exposed hydrophobic residues introduced during design were reverted to their WT identities.

### Screening of T=4 nanoparticles by co-purification

Three tricistronic genes were ordered from IDT. An N-terminal GFP was included on the A subunit of the pentasymmetron heterotrimer as a mass tag. A C-terminal 6x-histidine tag was added to the C subunit. Genes were expressed at medium scale (see above). Clarified lysate was loaded onto 1 mL of Ni-NTA resin (Thermo Fisher Scientific) pre-equilibrated in wash buffer. After washing with 3 column volumes of wash buffer, the protein was eluted with 2 column volumes of elution buffer. Eluate was screened for the presence of all three gene products by SDS-PAGE.

### Purification of co-expressed GI_4_-F7

GI_4_-F7 nanoparticles expressed tricistronically at large scale were purified by loading on a 5 mL HisTrap FF column (Cytiva) equilibrated in wash buffer (50 mM Tris pH 8.0, 250 mM NaCl, 20 mM imidazole, 1 mM DTT). After loading, the column was washed with 3-5 column volumes of wash buffer and protein was eluted with a gradient into 100% elution buffer (50 mM Tris pH 8.0, 250 mM NaCl, 500 mM imidazole, 1 mM DTT) over 40 minutes at 3 mL/min. The major fractions from elution were pooled, concentrated to ∼1 mL, and loaded onto an equilibrated Sephacryl S-500 HR 10/300 GL. SEC buffer was 25 mM Tris pH 8.0, 150 mM NaCl, 1 mM DTT.

### Purification of GI_4_-F7 heterotrimeric and homotrimeric components

For*in vitro* assembly, the heterotrimeric component of GI _4_-F7, comprising only the A and B chains, was expressed bicistronically. The A chain was modified with an N-terminal 6x-histidine tag. When expressed this way, some AAA nanoparticles and BBB homotrimers likely assemble in addition to AAB and ABB heterotrimers. To purify AAB from ABB heterotrimers, the bicistronic gene was expressed at large scale with the modification that 0.75% CHAPS was added to the lysis buffer and DTT was omitted. Clarified lysate was purified with a 5 mL HisTrap FF column as described above. Elution chromatograms contained three peaks. The first peak was predominantly the ABB heterotrimer, the second peak was predominantly AAB heterotrimer, and the third peak was predominantly AAA homotrimers assembled into an I3-01 like particle, A_60_. Any BBB homotrimer would be in the flow-through. The first and second peaks were pooled separately and concentrated to ∼1 mL. To remove any residual A_60_ we further purified the concentrated fractions on a Superose 6 Increase 10/300 column. The SEC buffer was 25 mM Tris pH 8.0, 150 mM NaCl, 0.75% w/v CHAPS. Glycerol was added to purified heterotrimer to a final concentration of 5%, the concentration was determined by A280, and 1 mL aliquots were flash-frozen in liquid nitrogen. Aliquots were stored at -80 °C until use. The homotrimer components were expressed at large scale and purified by IMAC in the same way as the co-expressed GI_4_-F7 nanoparticles except that 1% CHAPS was added to all buffers. It was further purified by SEC on a HiLoad 26/600 Superdex 200 PG column in 25 mM Tris pH 8.0, 150 mM NaCl, 5% Glycerol, 1.0% w/v CHAPS, 1 mM DTT. The total trimer protein concentration was measured by A280, flash-frozen in liquid nitrogen in 1 mL aliquots, and stored at -80 °C until use.

### *in vitro* assembly of GI_T_-F7 nanoparticles

To assemble GI_T_-F7 nanoparticles, components were mixed at various stoichiometries depending on the target assembly state, and dialyzed into 0% CHAPS overnight at room temperature in a 30 kDa cutoff dialysis cassette. Because the AAB and ABB heterotrimers do not directly interact with the BBB homotrimer, those components were mixed first, followed by addition of the CCC homotrimer. Nanoparticles were prepared fresh for each experiment, or stored at 4 °C for up to three days. To assemble BBB-CCC 2D arrays, the components were first individually dialyzed to remove CHAPS, and then mixed at a 1:1 stoichiometric ratio and allowed to assemble overnight at room temperature.

### Characterization of assemblies

Assemblies were characterized in solution by dynamic light scattering (DLS). Samples were measured using an UNcle (UNchained Labs) according to the manufacturer’s directions. Briefly, 8.8 μL of sample was loaded in triplicate into the capillary cassette. For each replicate, 10 acquisitions 10 seconds in length were collected. Assemblies were further characterized by negative-stain electron microscopy. Samples were diluted to between 0.1 and 0.5 mg/mL total protein depending on the assembly stoichiometry, applied to a glow-discharged thick carbon film 400 mesh copper grid (Electron Microscopy Sciences), and stained with 2% uranyl formate. Care was taken to ensure the stain thickness was sufficient to support the larger assemblies. Micrographs were collected on a Talos L120C (FEI) at up to 48,000× magnification. Individual micrographs were processed with ImageJ.

### CryoEM sample preparation, data collection and data processing

Three microliters of 3 mg/mL GI_4_-F7, GI_9_-F7, and GI_16_-F16 were loaded onto freshly glow discharged R 2/2 UltrAuFoil grids, prior to plunge freezing using a Vitrobot Mark IV (ThermoFisher Scientific) with a blot force of 0 and 6 sec blot time at 100% humidity and 22°C. Data were acquired using an FEI Titan Krios transmission electron microscope operated at 300 kV and equipped with a Gatan K3 direct detector and Gatan Quantum GIF energy filter, operated in zero-loss mode with a slit width of 20 eV. For GI_4_-F7 and GI_9_-F7, automated data collection was carried out using Leginon ^92^ at a nominal magnification of 105,000× with a pixel size of 0.843 Å. 7,249 and 2,558 micrographs were collected with a defocus range comprised between -0.5 and -2.5μm, respectively. The dose rate was adjusted to 15 counts/pixel/s, and each movie was acquired in super-resolution mode fractionated in 75 frames of 40 ms. For the GI_16_-F7 data set, automated data collection was carried out using Leginon ^92^ at a nominal magnification of 64,000× with a pixel size of 1.42 Å. 2,268 micrographs were collected with a defocus range between -0.5 and -3.5μm. The dose rate was adjusted to 15 counts/pixel/s, and each movie was acquired in super-resolution mode fractionated in 50 frames of 100 ms. Movie frame alignment, estimation of the microscope contrast-transfer function parameters, particle picking, and extraction were carried out using Warp ^93^.

Two rounds of reference-free 2D classification were performed using CryoSPARC ^94^ to select well-defined particle images. These selected particles were subjected to two rounds of 3D classification with 50 iterations each (angular sampling 7.5° for 25 iterations and 1.8° with local search for 25 iterations) using Relion ^95^with an initial model generated with ab-initio reconstruction in cryoSPARC. 3D refinements were carried out using non-uniform refinement along with per-particle defocus refinement in CryoSPARC. Selected particle images were subjected to the Bayesian polishing procedure ^96^ implemented in Relion 3.1 before performing another round of non-uniform refinement in cryoSPARC followed by per-particle defocus refinement and again non-uniform refinement. To further improve the density of the ASU, the particles were symmetry-expanded and subjected to focus 3D classification without refining angles and shifts. Particles belonging to classes with the best resolved ASU density were selected and then subjected to local refinement using CryoSPARC. Local resolution estimation,and sharpening were carried out using CryoSPARC. Reported resolutions are based on the gold-standard Fourier shell correlation (FSC) of 0.143 criterion and Fourier shell correlation curves were corrected for the effects of soft masking by high-resolution noise substitution ^97,98^.

### Model building and refinement

UCSF Chimera ^99^ and Coot ^100^ were used to fit atomic models into the cryoEM maps. GI_4_-F7 and GI_9_-F7 asu models were refined and relaxed using Rosetta using sharpened and unsharpened maps ^101,102^. For GI_4_-F7 or GI_9_-F7 icosahedral model, all of the side chains of GI_4_-F7 or GI_9_-F7 asu model are truncated except Gly, Cys and Pro residues and the symmetry related copies were generated in ChimeraX with Cryo EM maps.

### Alignments and images

To align the cryoEM models to the design model, both models were centered at the origin and their icosahedral symmetry axes aligned in PyMOL ^103^. The CαRMSD was calculated using the “rms_cur” function in PyMOL. To measure deviations in the rigid body degrees of freedom, copies of the pentasymmetron, disymmetron, and trimer (or trimers for GI_9_-F7) from the cryoEM model were aligned to the design model using the “super” function in PyMOL. We then calculated the rotations and translations from the transformation matrix between the corresponding component of the original cryoEM model and the aligned cryoEM model. We applied the same approach to the heterotrimer (and homotrimer for GI_9_-F7) components to obtain rotations and translations within the pentasymmetron, disymmetron, and homotrimer components, respectively. We found that the “super” function in PyMOL was very sensitive to chain and residue numbering, as well as some of the minor differences between the design model and cryoEM model. Therefore, for all alignments using PyMOL, we made sure to harmonize residue numbering, chain IDs, and remove any residues present in only one model or the other. For that reason, aligned images were generated using the “mm” command in ChimeraX ^104^ and verified to ensure that the alignments closely matched those generated on the trimmed models created with the “super” function in PyMOL.

### Scripts and plots

All data was processed and plotted using python 3.8.8, Matplotlib 3.3.4, and seaborn 0.11.1.

