## Supplemental Data for "Hierarchical design of pseudosymmetric protein nanoparticles"

### Supplemental Figures

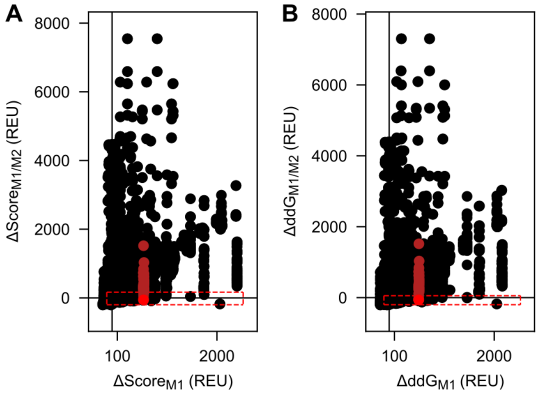


**Fig. S1: Rosetta metrics for all single and pairwise 1WA3 trimer interface mutants.** (**A**) ΔddG filter metric. (**B**) ΔScore metric. Dark red points correspond to the single mutation P114F. The bright red point corresponds to the double mutant P114F/F131V. The red dotted boxes represent cutoffs used to select mutants for testing.

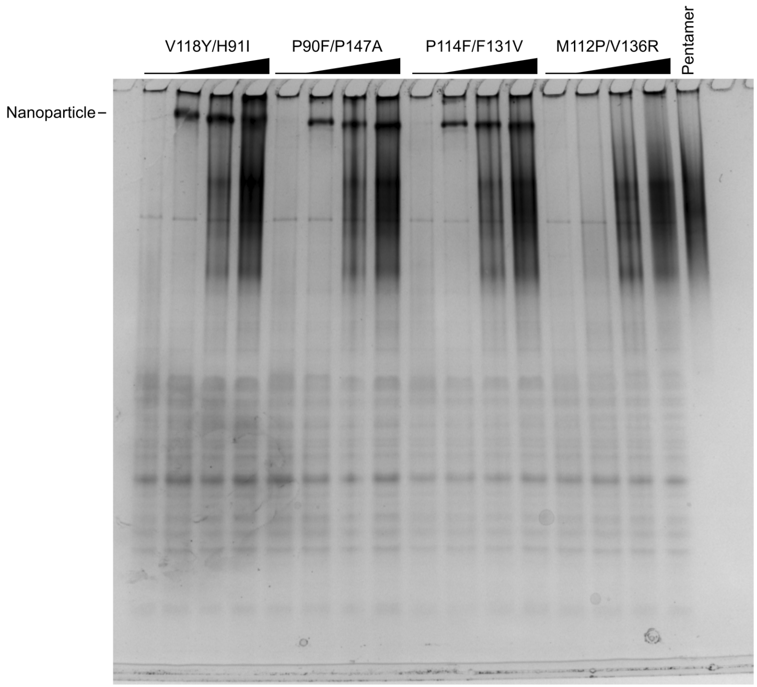


**Fig. S2: Native-PAGE screen of double mutants.** Recovery of trimer geometry was assayed by assembling double mutant I53-50A trimers in clarified *E. coli* lysates with purified I53-50B pentamer and evaluating the presence or absence of I53-50 nanoparticles by native PAGE. Black wedges indicate increasing pentamer concentration in each series of assembly reactions.

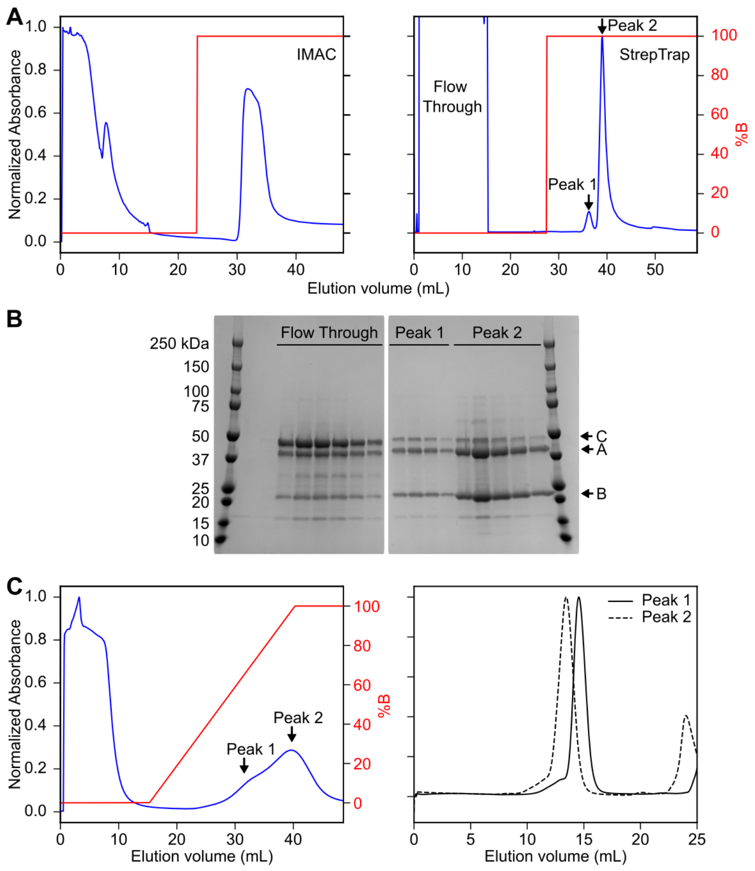


**Fig. S3: Purification and characterization of "ABC" tricistronic and "AB" bicistronic constructs.** (**A**) The ABC heterotrimer was purified by (*left*) IMAC with a step elution followed by (*right*) StrepTrap purification. The A chain contained a hexa-histidine and SUMO tag, the B chain contained a Strep tag, and the C chain contained sfGFP and avi tags. The eluate of this two-step purification method should therefore only contain trimers that include both the A and B chains. An optimal result would be equimolar amounts of the A, B, and C chains. (**B**) SDS-PAGE of the StrepTrap purification revealed that the eluate contained an excess of the A and B chains and less of the C chain. To test the ability of the A and B chains only to assemble into heterotrimers, we expressed an AB bicistronic gene and (**C**) purified the resulting proteins by (*left*) IMAC with a gradient elution. Two broad and overlapping peaks were observed. The leading half of the first peak and trailing half of the second peak were collected and (*right*) further purified by SEC. Peak 2 has a lower retention volume than peak 1, suggesting a difference in molecular weight. These results are consistent with assembly of an ABB heterotrimer (earlier IMAC elution, later SEC elution) and an AAB heterotrimer (later IMAC elution, earlier SEC elution). We confirmed this interpretation by native mass spectrometry (**Fig. 1G**)

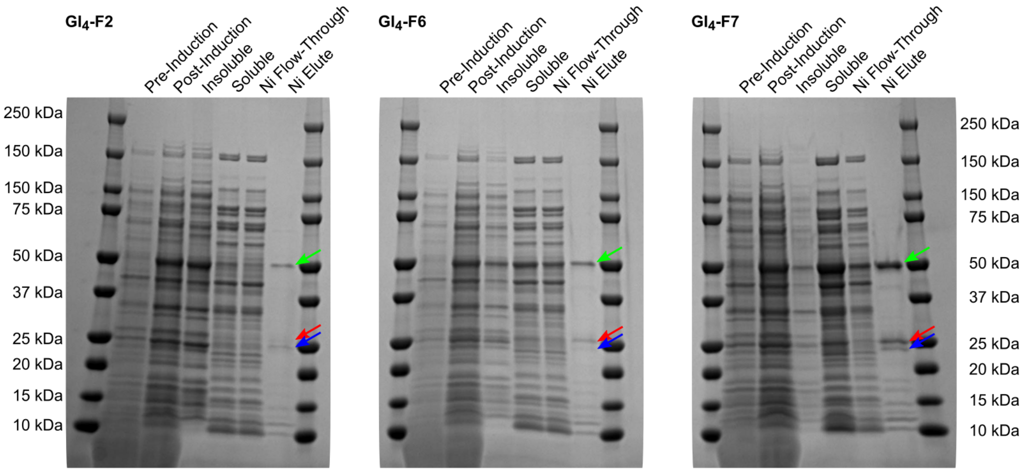


**Fig. S4: SDS-PAGE of GI$_4$ designs by Ni$^{2+}$ pull-down assay.** Expression and screening by SDS-PAGE for GI$_4$ designs. Bands for chains A (green arrow), B (blue arrow), and C (red) arrow are indicated. The presence of all three bands in the Ni Elute lanes of GI$_4$-F6 and GI$_4$-F7 indicates interactions between the A, B, and C chains.

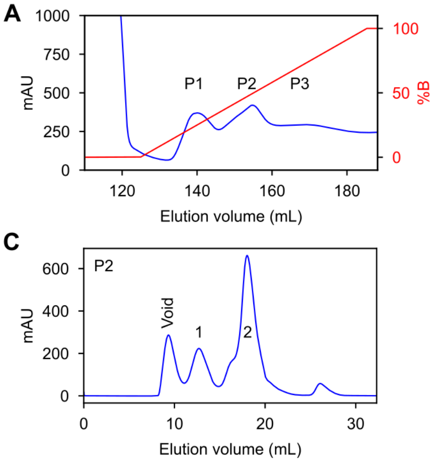


**Fig. S5: Purification of heterotrimer components from $A_{60}$.** (**A**) HisTrap elution chromatogram. Blue, absorbance at 280 nm; red, gradient elution. Peak 1 (P1) is predominantly ABB, P2 is predominantly AAB, and P3 is predominantly the A chain, which assembles into 60-subunit I3-01-like nanoparticles. (**B**) Superdex 200 Increase 10/300 chromatogram of P1 from the HisTrap elution. The first peak following the void volume (1) is predominantly I3-01-like nanoparticles and (2) is predominantly ABB heterotrimer. (**C**) Superdex 200 Increase 10/300 chromatogram of P2 from the HisTrap elution. (1) is predominantly I3-01-like nanoparticles and (2) is predominantly AAB heterotrimer. (**D**) Superdex 200 Increase 10/300 chromatogram of P3 from the HisTrap elution. (1) is predominantly I3-01-like nanoparticles and (2) is predominantly AAB heterotrimer.

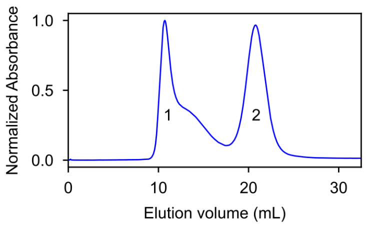


**Fig. S6: Purification of $GI_4$-F7 by SEC.** SEC purification of $GI_4$-F7 on a Sephacryl S-500 HR 10/300 GL column. Peak 1 contains the assembly while peak 2 is residual homotrimer component.

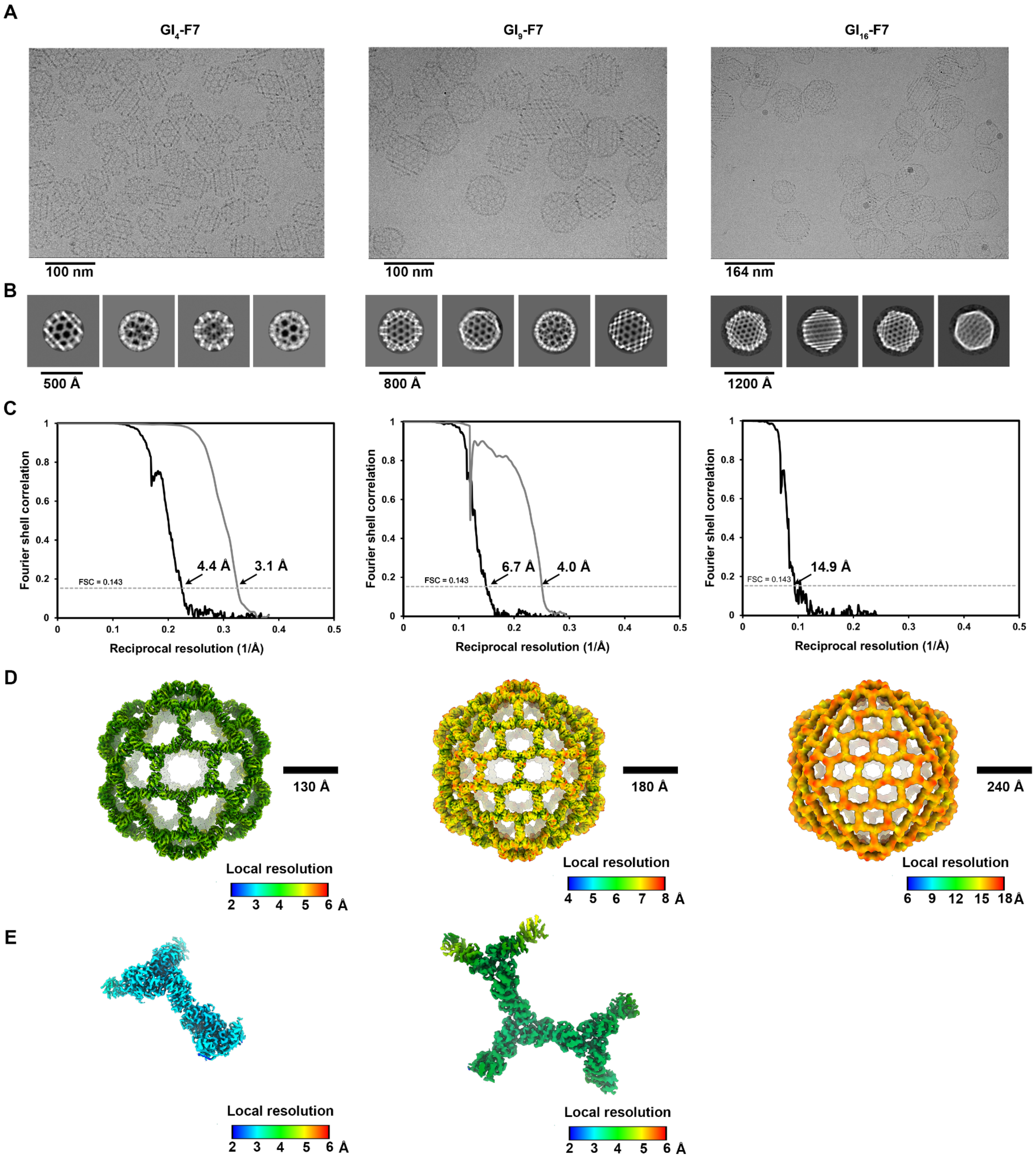


**Fig. S7: CryoEM data processing.** (**A-B**) Representative electron micrographs (A) and 2D class averages (B) of $GI_4$-F7 (left), $GI_9$-F7, (middle) and $GI_{16}$-F7 (right). (**C**) Gold-standard Fourier shell correlation curves for the 3D reconstructions of $GI_4$-F7 (left), $GI_9$-F7 (middle) and $GI_{16}$-F7 (right) (black line) and locally refined asus (gray lines). (**D-E**) Local resolution maps calculated using cryoSPARC for (D) the 3D reconstructions of $GI_4$-F7 (left), $GI_9$-F7 (middle), and $GI_{16}$-F7 (right) as well as (E) the locally refined ASUs of $GI_4$-F7 (left) and $GI_9$-F7 (middle). The 0.143 cutoff is indicated by a horizontal dashed line.

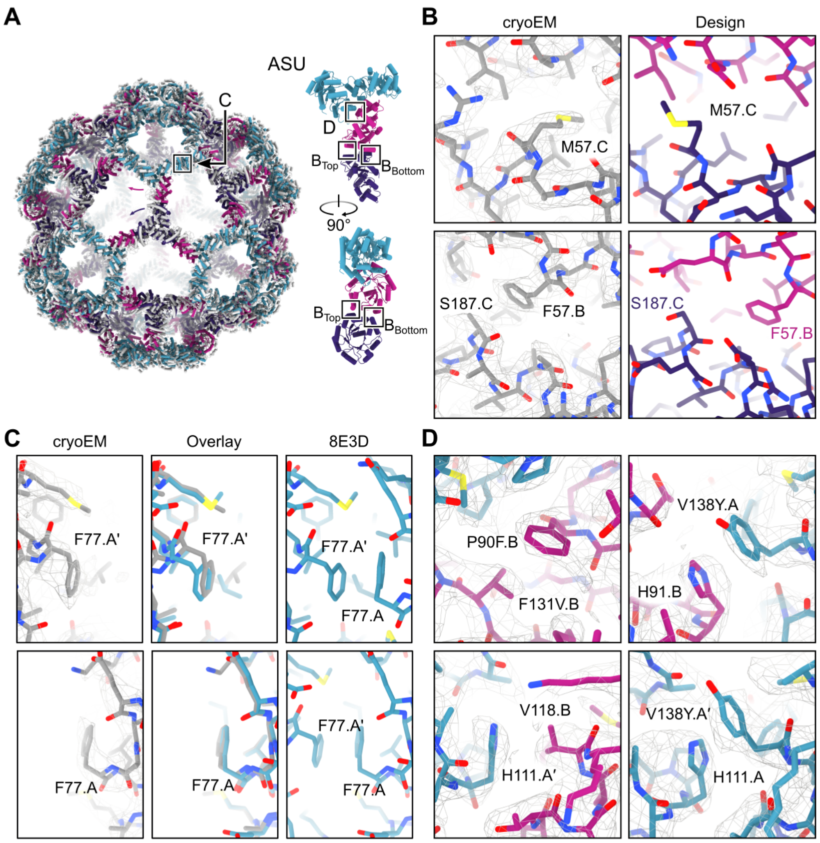


**Fig. S8: Structural details of GI$_4$-F7.** (**A**) Alignment of the complete cryoEM model to the design model. Major rigid-body DoF deviations are indicated with arrows. Two views of the ASU are shown. Approximate locations of each inset (B, C, and D) are indicated. (**B**) Comparison between the cryoEM model (left) and design model (right) of the newly designed nanoparticle (B-C) interface. *Top row*, M57 on the CCC-homotrimer changes rotamer to occupy a void in the interface in the design model. *Bottom row*, F57 on the B chain of the AAB heterotrimer packs against S187 of the CCC homotrimer in the cryoEM model, instead of A190 in the CCC homotrimer as in the design model. (**C**) Comparison of the I3-01 (A-A) interface observed in the cryoEM model to a previously published structure (PDB ID 8ED3). *Top row*, slight rigid-body deviations from perfect two-fold symmetry in one copy of the A chain. *Bottom row,* very little deviation from perfect two-fold symmetry (**D**) Details of the density maps in the regions of the pseudosymmetry-generating mutations within the AAB heterotrimer interface.

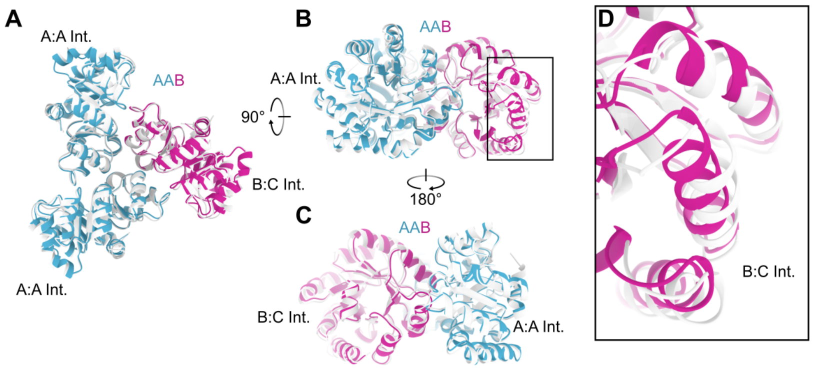


**Fig. S9: Comparison of the AAB heterotrimer design model to the cryoEM model from $GI_4$-F7.** Alignment of the AAB heterotrimer cryoEM model to the design model is viewed (**A**) from the top, towards the center of the nanoparticle along the three-fold symmetry axis; (**B**) from the side, tangential to the nanoparticle surface; and (**C**) from the other side, tangential to the nanoparticle surface. The position of the A:A and newly designed B:C interfaces are indicated. (**D**) Detail of the B side of the B:C interface, highlighting the most significant deviations from the design model.

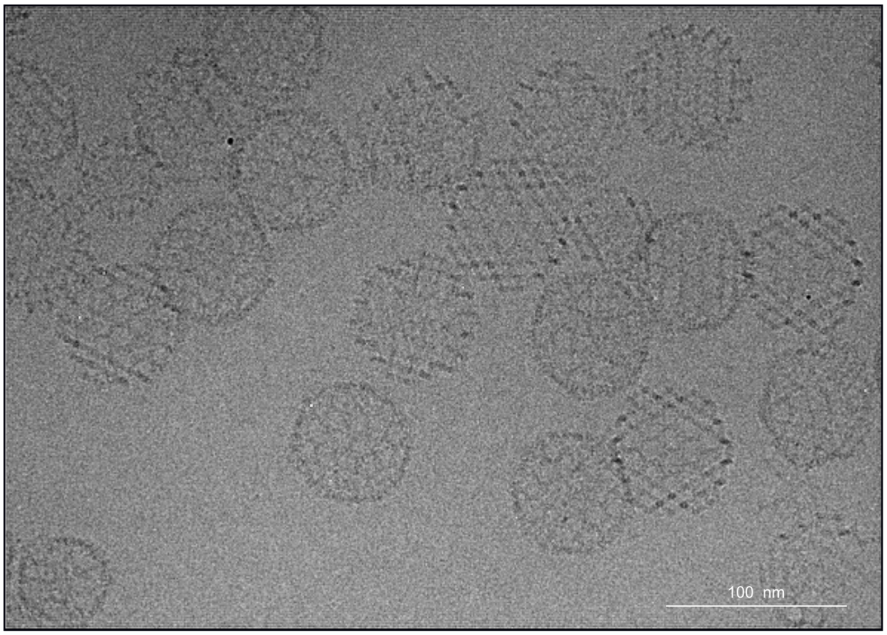


**Fig. S10: Field view micrograph of $GI_9$-F7.** CryoEM field view micrograph of samples enriched for $GI_9$-F7 by SEC purification. Both $GI_9$-F7 (large particles) and $GI_4$-F7 (e.g., bottom-left corner) are clearly visible.

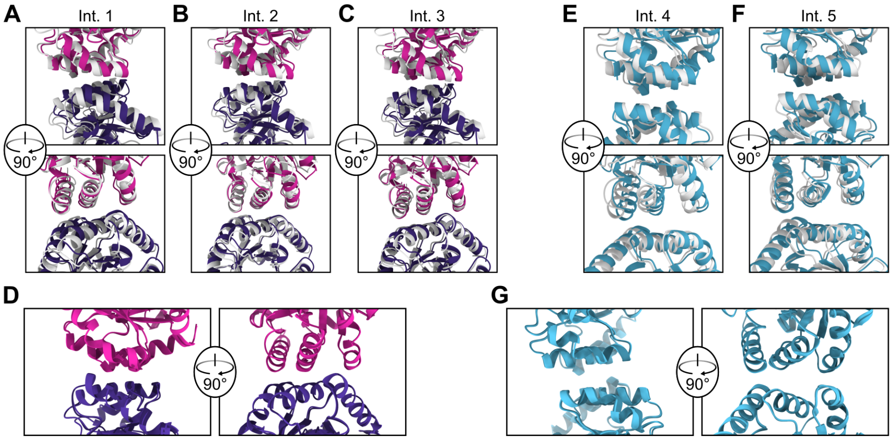


**Fig. S11: Structural details of $GI_9$-F7.** (**A-C**) Alignment of the $GI_9$-F7 design model chain B (magenta) and chain C (purple) protein-protein interface to the corresponding chains of the cryoEM model (gray). Each of the three interfaces between B and C chains in the ASU are shown. (**D**) The protein-protein interface between chain B and C from the cryoEM model of $GI_4$-F7 (light colors) aligned to the same interface from the cryoEM model of $GI_9$-F7 (dark colors). (**E**) Alignment of design model to the cryoEM model for the I3-01 interface in the pentasymmetron and (**F**) disymmetron. (**G**) Alignment of the I3-01 interface from the cryoEM models of $GI_4$-F7 (light blue) and $GI_9$-F7 (dark blue).

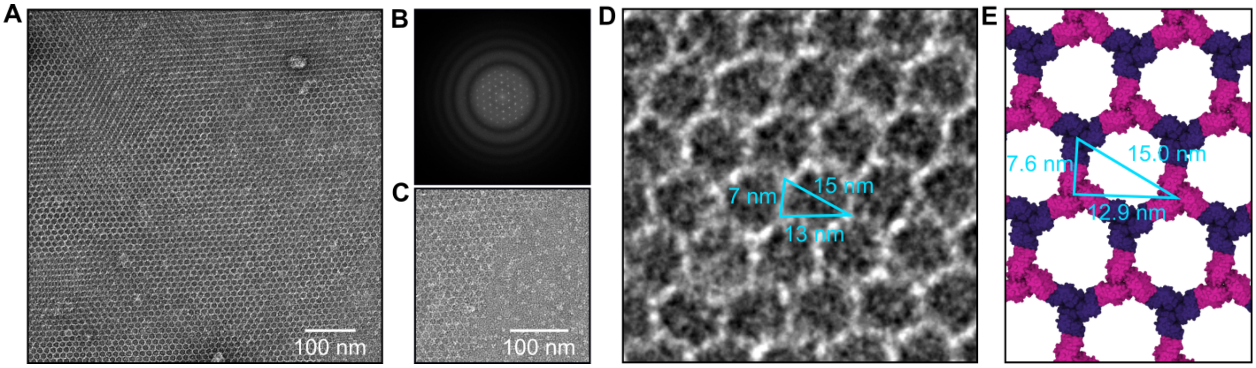


**Fig. S12: Hexagonal 2D array characterization by negative stain EM.** (**A**) An example of the regular hexagonal array formed by mixing BBB and CCC homotrimers by negative stain EM. (**B**) Power spectrum of the micrograph shown in panel A, confirming the periodic arrangement of the array. (**C**) The edge of the array is jagged, with free trimeric components visible. (**D**) Measurement of the array dimensions are consistent with (**E**) the design model.

### Supplemental Tables

**Supplementary Table S1: Table of all single and double mutants and outcome.**

| Single mutant | Assembles in lysate? | Single mutant | Assembles in lysate? | Single mutant | Assembles in lysate? |
|---|---|---|---|---|---|
| R17Y | Yes | T113V | Yes | **F140K** | **No** |
| T69E | Yes | P114Q | Yes | F140H | Yes |
| T69G | Yes | P114S | Yes | F140W | Yes |
| T71A | Yes | **P114F** | **No** | A143D | Yes |
| T71D | Yes | T115K | Yes | A143E | Yes |
| T71E | Yes | T116C | Yes | A143Y | Yes |
| T71K | Yes | T116D | Yes | P147A | Yes |
| T71L | Yes | T116G | Yes | P147D | Yes |
| T71N | Yes | T116L | Yes | P147H | Yes |
| T71Y | Yes | T116M | Yes | P147M | Yes |
| P90E | Yes | T116N | Yes | P147R | Yes** |
| **P90F** | **No** | T116Q | Yes | P147S | Yes |
| P90K | Yes | T116S | Yes | F148C | Yes |
| P90Q | Yes | T116T | Yes | F148E | Yes |
| P90R | Yes | **T116W*** | **No** | F148H | Yes |
| H91C | Yes | V118H | Yes | F148Q | Yes |
| H91D | Yes | V118K | Yes | F148Y | Yes |
| H91F | Yes | V118M | Yes | | |
| H91G | Yes | V118N | Yes | | |
| H91I | Yes | V118S | Yes | | |
| H91S | Yes | V118W | Yes | | |
| D93R | Yes | **V118Y** | **No** | | |
| **M112P** | **No** | K122D | Yes | | |
| M112Q | Yes | K122E | Yes | | |
| M112V | Yes | F131E | Yes** | | |
| T113D | Yes | E134G | Yes | | |
| T113F | Yes | E134K | Yes | | |
| T113I | Yes | V135P | Yes | | |
| T113L | Yes | V136R | Yes | | |
| T113M | Yes | V136Y | Yes | | |
| T113P | Yes | V136W | Yes | | |
| T113Q | Yes | Q139E | Yes | | |

| Double mutant | Assembles in lysate? |
|---|---|
| M112P/V136R | No |
| P90F/P147A | Yes |
| P114F/F131V | Yes |
| V118Y/H91I | Yes |

* Mutant at three-fold position required screening in a tricistronic gene construct and was not pursued further.
** Did not assemble in one replicate or had a weak assembly band.

**Supplementary Table S2. Amino acid sequences for novel proteins used in this study.**

>Pseudosymmetric I53-50A A Chain
MGHHHHHHHHHHGSLQDSEVNQEAKPEVKPEVKPETHINLKVSDGSSEIFFKIKKTTPLRRLMEAFAKRQGKEMDSLRF
LYDGIRIQADQAPEDLDMEDNDIIEAHREQIGGSEKAAKAEEAARKMEELFKKHKIVAVLRANSVEEAIEKAVAVFAGGVHL
IEITFTVPDADTVIKALSVLKEKGAIIGAGTVTSVEQCRKAVESGAEFIVSPHLDEEISQFCKEKGVFYMPGVMTPTELYKA
MKLGHDILKLFPGEVVGPQFVKAMKGPFPNVKFVPTGGVNLDNVCKWFKAGVLAVGVGKALVKGKPDEVREKAKKFVK
KIRGCTE

>Pseudosymmetric I53-50A B Chain
MKMEELFKEHKIVAVLRANSVEEAISKALAVFAGGVHLIEITFTVPDADQVIKELEFLKEAGAIIGAGTVTSVEQCREAVESG
AEFIVSFHLDEEISQFCKEEGVFYMPGVMTPTELVKAMKLGHTILKL**V**PGEVVGPQFVEAMKGPFPNVKFVPTGGVNLD
NVCEWFEAGVLAVGVGSALVEGEPAEVAELAIRFVEKIRGCTEGSGWSHPQFEK

>Pseudosymmetric I53-50A C Chain
MKGEELFTGVVPILVELDGDVNGHKFSVRGEGEGDATNGKLTLKFICTTGKLPVPWPTLVTTLTYGVQCFARYPDHMKQ
HDFFKSAMPEGYVQERTISFKDDGTYKTRAEVKFEGDTLVNRIELKGIDFKEDGNILGHKLEYNFNSHNVYITADKQKNGI
KANFKIRHNVEDGSVQLADHYQQNTPIGDGPVLLPDNHYLSTQSVLSKDPNEKRDHMVLLEFVTAAGITHGMDELYKGG
SGGSGGKMEELFKKHKIVAVLRANSVEEAIEKAVAVFAGGVHLIEITFTVPDADTVIKALSVLKEKGAIIGAGTVTSVEQCRK
AVESGAEFIVSPILDEEISQFCKEKGVFYMPGVMTFTELVKAMKLGHTILKLFPGEVVGPQFVKAMKGAFPNVKFVPTGG
VNLDNVCEWFKAGVLAVGVGSALVKGTPDEVREKAKAFVEKIRGCTEGSGLNDIFEAQKIEWHE

>Pseudosymmetric $GI_T$ A Chain
MGSHHHHHHGSEKAAKAEEAARKMEELFKEHKIVAVLRANSVEEAKKKALAVFLGGVHLIEITFTVPDADTVIKELSFLKE
MGAIIGAGTVTSVEQCREAVESGAEFIVSPHLDEEISQFCKEEGVFYMPGVMTPTELYKAMKLGHTILKLFPGEVVGPQF
VEAMKGPFPNVKFVPTGGVNLDNVCEWFEAGVLAVGVGSALVEGTPVEVAEKAKAFVEKIEGCTE

>Pseudosymmetric $GI_T$ B Chain
MKMEELFKEHKIVAVLRANSVEEAISKALAVFAGGVHLIEITFTVPDADQVIKELEFLKEAGAIIGAGTVTSVEQCREAVESG
AEFIVSFHLDEEISQFCKEEGVFYMPGVMTPTELVKAMKLGHTILKL**V**PGEVVGPQFVEAMKGPFPNVKFVPTGGVNLD
NVCEWFEAGVLAVGVGSALVEGEPAEVAELAIRFVEKIRGCTE

>Homotrimeric $GI_T$ BBB
MKMEELFKEHKIVAVLRANSVEEAISKALAVFAGGVHLIEITFTVPDADQVIKELEFLKEAGAIIGAGTVTSVEQCREAVESG
AEFIVSPHLDEEISQFCKEEGVFYMPGVMTPTELVKAMKLGHTILKLFPGEVVGPQFVEAMKGPFPNVKFVPTGGVNLD
NVCEWFEAGVLAVGVGSALVEGEPAEVAELAIRFVEKIRGCTELEHHHHHH

>Homotrimeric $GI_T$ CCC
MKMEELFKEHKIVAVLRANSREEAIEIALAVFAGGVHLIEITFTVPDADEVIKRLEMLKRAGAIIGAGTVTSVEQCREAVESG
AEFIVSPHLDEEISQFCKEEGVFYMPGVMTPTELVKAMKLGHTILKLFPGEVVGPQFVEAMKGPFPNVKFVPTGGVNLD
NVCEWFEAGVLAVGVGSALVEGKPSEVAEKARRFVKKIRGCTEGSLEHHHHHH

Appended sequences including SUMO, GFP, and deca- or hexa-histidine, avi- and strep- tags are underlined.
Pseudosymmetrizing mutations are highlighted.

**Supplementary Table S3. CryoEM data collection and refinement statistics.**

| | $GI_4$-F7<br><br>EMD XXXX | $GI_4$-F7<br>(local refinement)<br>PDB XXXX<br>EMD XXXX | $GI_9$-F7<br><br>EMD XXXX | $GI_9$-F7<br>(local refinement)<br>PDB XXXX<br>EMD XXXX | $GI_{16}$-F7<br><br>EMD XXXX |
|---|---|---|---|---|---|
| **Data collection and processing** | | | | | |
| Magnification | 105,000 | 105,000 | 105,000 | 105,000 | 64,000 |
| Voltage (kV) | 300 | 300 | 300 | 300 | 300 |
| Electron exposure ($e^{-}$/Å$^2$) | 60 | 60 | 60 | 60 | 37 |
| Defocus range (μm) | -0.5 - -2.5 | -0.5 - -2.5 | -0.5 - -2.5 | -0.5 - -2.5 | -0.5 - -3.5 |
| Pixel size (Å) | 0.843 | 0.843 | 0.843 | 0.843 | 1.42 |
| Symmetry imposed | I | C1 | I | C1 | I |
| Initial particle images (no.) | 154,574 | | 18,611 | | 1,226 |
| Final particle images (no.) | 120,979 | 984,020 | 1,956 | 795,360 | 1,083 |
| Map resolution (Å) | 4.4 | 3.1 | 6.7 | 4.0 | 14.9 |
| FSC threshold | 0.143 | 0.143 | 0.143 | 0.143 | 0.143 |
| **Refinement** | | | | | |
| Map resolution (Å)<br>FSC threshold | | 0.143 | | 0.143 | |
| Map sharpening *B*factor (Å$^2$) | -269 | -143 | -530 | -138 | |
| Model composition<br>Non-hydrogen atoms<br>Protein residues<br>Ligands | | 5687<br>813<br>0 | | 9161<br>1827<br>0 | |
| B factors (Å2)<br>Protein<br>Ligand | | 15.53 | | 73.75 | |
| **Validation** | | | | | |
| MolProbity score | | 1.9 | | 1.06 | |
| Clashscore | | 5.86 | | 0.71 | |
| Poor rotamers (%) | | 2.94 | | 0.13 | |
| Ramachandran plot | | | | | |
| Favored (%) | | 96.52 | | 98.95 | |
| Allowed (%) | | 3.11 | | 0.5 | |
| Disallowed (%) | | 0.37 | | 0.55 | |

**Supplementary Table S4: Deviations observed in the cryoEM reconstruction of $GI_4$-F7 compared to the design model.**

| Component | Axis | Translation (Å) | Rotation (Degrees) |
|---|---|---|---|
| Pentasymmetron | 5-fold | 5.8 | 5.9 |
| AAB-Heterotrimer | // Local 3-fold | 1.1 | 1.3 |
| | ⊥ Local 3-fold | 1.6 | – |
| | AAB:AAB Interface | – | 1.6, -2.8 |
| | AAB:CCC Interface | – | -0.2 |
| Trimer | 3-fold | 4.0 | 12.4 |

**Supplementary Table S5: Deviations observed in the cryoEM reconstruction of $GI_9$-F7 compared to the design model.**

| Component | Axis | Translation (Å) | Rotation (Degrees) |
|---|---|---|---|
| Pentasymmetron AAB heterotrimer | Icosahedral 5-fold | 7.7 | 7.2 |
| | ∥* Local 3-fold | 1.3 | 1.8 |
| | ⊥ Local 3-fold | 1.9 | -- |
| | AAB:AAB Interface | -- | 1.9, -3.6 |
| | AAB:CCC Interface | -- | 0.3 |
| Disymmetron ABB heterotrimer | Icosahedral two-fold | 10.0 | 8.2 |
| | ∥ Local 3-fold | 0.4 | 2.3 |
| | ⊥ Local 3-fold | 1.3 | -- |
| | ABB:ABB Interface | -- | -1.6 |
| | ABB:CCC Interface | -- | -0.8, 3.7 |
| 3× Trimer CCC homotrimer | Icosahedral 3-fold | 6.5 | 5.4 |
| | ∥ Local 3-fold | 0.0 | -6.5 |
| | ⊥ Local 3-fold | 1.23 | -- |
| | CCC:ABB Interface | -- | 0.3, -1.8 |
| | CCC:AAB Interface | -- | -0.6* |

*The symbol ∥ indicates deviations parallel to the indicated symmetry axis, ⊥ indicates deviations perpendicular to the indicated axis.

**Supplementary Table S6: DLS results obtained from assembly reactions corresponding to T numbers 4 to 100.**

| T | Mean Z-Ave. dia. (nm) | StdDev Z-Ave. dia. (nm) | PDI |
|---|---|---|---|
| 4 | 47.5 | 0.4 | 0.048 |
| 9 | 69.9 | 0.5 | 0.154 |
| 16 | 95 | 1 | 0.121 |
| 25 | 113.2 | 0.5 | 0.127 |
| 36 | 129 | 2 | 0.134 |
| 49 | 148 | 2 | 0.136 |
| 64 | 162 | 1 | 0.136 |
| 81 | 174 | 1 | 0.126 |
| 100 | 189 | 1 | 0.133 |
